# Brain responses to predictable structure in auditory sequences: From complex regular patterns to tone repetition

**DOI:** 10.1101/2024.07.18.604117

**Authors:** Rosy Southwell, Candida Tufo, Maria Chait

## Abstract

In classical studies of auditory pattern learning, neuronal responses exhibit repetition suppression, where sequences of repeated tones show a reduced evoked response. This may be due in part to adaptation, but is also hypothesized to indicate suppression of expected stimuli. Repetition suppression is thought to form a building block of regularity learning, and is the paradigmatic example of predictive coding in humans and other animals.

However, stimuli with more complex patterns appear to show the opposite effect. Predictable regular (REG) patterns distributed over a range of frequencies show a strongly enhanced sustained brain response compared to frequency-matched random (RAND) sequences, sitting at odds with the usual reduced evoked responses to predictable stimuli. A limiting factor in reconciling these findings is that they are obtained using different stimuli and analysis methods.

This human EEG study (N=20) brings together auditory sequence predictability and repetition in a single paradigm, based on rapidly unfolding tone pip patterns, incorporating sequences consisting of exactly repeating tones at a single frequency, alongside REG and RAND of varying complexity. We demonstrate that regularity is associated with increased sustained responses, offset responses, tone locked responses and cycle-locked responses. We further show that both repetition suppression and repetition enhancement occur over different timescales, and that sustained brain responses to simple repetition show qualitatively different effects than to more complex regularities during automatic tracking of stimulus statistics. Our results indicate a system for automatic monitoring of predictability in the auditory environment, which is distinct from, but concurrent with, repetition suppression.

**NEW & NOTEWORTHY:** We investigate how human brain response patterns evolve as auditory patterns become progressively simplified, focusing on the impact of sequence structure on various aspects of EEG responses, both evoked and induced. By contrasting EEG responses across different phases of the stimulus—onset, sustained, and offset—we uncover the interplay between repetition suppression and predictability. This approach allows us to discern how the brain processes and adapts to changing regularities in sensory input.

## INTRODUCTION

Detecting regular structure in sensory input is a prerequisite for forming a predictive model of the world, thought to be fundamental to brain function (Friston, 2005; Bendixen, 2014; de Lange et al, 2018; Heilbron & Chait, 2018). Recent work in the auditory modality has revealed that these processes can be tracked by measuring the M/EEG sustained response evoked by unfolding sound sequences (Barascud et al, 2016; Southwell et al, 2017; Southwell & Chait 2018; Hermann et al, 2018; 2019; 2022; 2023; Al Jaja et al, 2020; Hu et al, 2024; Zhao et al, 2024). Studies utilizing patterns of tone pips arranged in regular repeating sequences (REG) or presented randomly (RAND) have consistently shown that the sustained response to REG patterns underpinned by a network of auditory cortical, inferior frontal gyrus, and hippocampal sources, surpasses that of RAND controls. The latency at which REG responses diverge from RAND, indicative of the timing of regularity detection, has been shown to be consistent with predictions from ideal observer models (Barascud et al, 2016; Skerritt-Davis & Elhilali, 2019; Hu et al, 2024), revealing an efficient automatic regularity extraction process. This has led to the hypothesis that the sustained response reflects the brain’s tracking of predictability, or precision (inverse variance) of the inferred predictive distribution of the sequence as it unfolds (Barascud et al, 2016; Zhao et al, 2024). The majority of research in this comparatively new area has focused on relatively long tone patterns, such as cycles of 10 repeating tones, aiming to probe complex processing while minimizing the influence of low-level factors such as adaptation (Heilbron & Chait, 2018).

Historically, the classic approach for studying sensitivity to auditory regularity has utilised very simple sound patterns (‘oddball paradigm’). This involves presenting long sequences of repeated tones, known as “standards,” which are occasionally interrupted by “deviant” tones (e.g., Näätänen et al., 1978; Garrido et al., 2008). A common observation is the reduction of evoked responses to the ‘standards’ (Baldeweg, 2006). Repetition suppression is often cited as evidence for predictive coding (Baldeweg, 2006; Garrido et al., 2009; Auksztulewicz and Friston, 2016), thought to result from top-down predictions suppressing responses to expected sensory input. However, using stimulus repetition as a model of predictability presents potential drawbacks. Firstly, simple repetition represents a very rudimentary pattern that may not fully capture the complexities of naturalistic sensory patterns. Moreover, the observed effects in such paradigms may be confounded by neuronal fatigue rather than solely reflecting top-down predictions informed by an internal representation of regularity (Nelken, 2014; though see e.g., Summerfield et al., 2008). To address these concerns, alternative procedures have been devised to disentangle the effects of repetition and expectation on auditory responses to repeated tones (Todorovic et al., 2011; Todorovic and de Lange, 2012). These studies have demonstrated that, alongside “repetition suppression,” “expectation suppression”—the reduction of responses to predicted tones—also shapes responses even in very simple tone sequences.

Interestingly the observations from these paradigms appear to contrast with those from the sustained response literature (above) where predictable patterns are associated with increased brain responses. By employing stimuli that allow for the dissociation of responses to individual tones and the sustained response, Hu et al. (2024) demonstrated that both effects can be measured concurrently in the unfolding brain response: sustained responses exhibited an increase to regular (REG) relative to random (RAND) patterns, consistent with the coding of predictability; responses to individual tones within the regular context exhibited reduced activity, consistent with findings in the “expectation suppression” literature. These multiplexed effects of predictability highlight the need to consider various experimental measures to gain a comprehensive understanding of how the brain responds to auditory patterns.

In this study, we investigate how and whether the sustained response changes in sequences with increasingly simple regularities. We utilize REG and RAND sequences along a continuum ranging from simple repetition to more complex wideband patterns (see Figure 1). REG1, consisting of one unique tone, represents simple repetition commonly used in literature demonstrating stimulus-specific adaptation, repetition suppression, and adaptation of the standard. REG3 is a pattern of three tones drawn from a pool of 20 values, while REG5 consists of five randomly selected tones forming a repeating pattern. We anticipate that as regularities become increasingly simple, measures reflecting the brain’s tracking of predictability will also reveal how these brain responses are susceptible to repetition suppression/adaptation effects.

**Figure 1:**
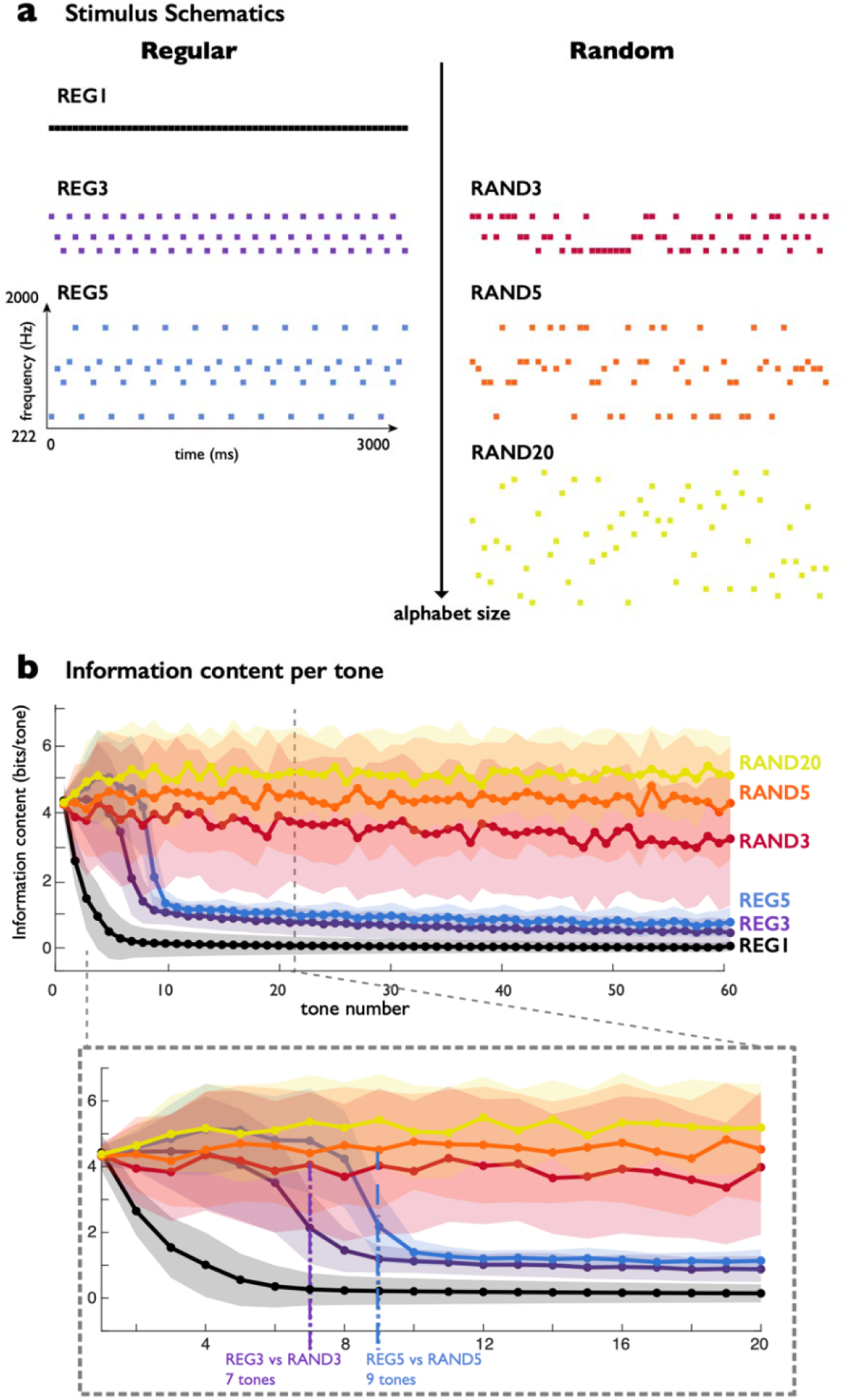
Stimuli. Tone-pip sequences varied in alphabet size (alph) from 1 to 20, and were either regular (REG) or random (RAND). a: Schematics depicting typical frequency patterns for the six conditions; squares represent individual tone-pips of duration 50 ms. For alph = 3 and 5, REG and RAND were generated in matched pairs with the same frequency content. ¬¬¬b: Mean information content per tone, estimated from a model of expectation computed over an entire experiment session. Mean information content computed over 108 trials of each condition. Shaded regions around lines represent standard deviation over trials. Bottom plot shows the same data re-plotted with a zoomed-in scale to show divergence between information content at the start of the sequence as regular pattern emerges.

Our predictions regarding the effect of predictability are anchored in an ideal observer model (IDyOM, see Figure 1b), previously employed to explain various phenomena related to neural and behavioural tracking of patterns in sound sequences (Pearce, 2005; Barascud et al., 2016; Bianco et al., 2020; Harrison et al., 2021). As depicted in Figure 1b, the information content (IC), a measure of predictability/surprise associated with each tone, drops early and sharply for REG1, indicating rapid detection of predictable structure. The discovery of regularity in REG3 and REG5, manifested as the divergence of IC between REG and RAND conditions, occurs after 7 and 9 tones, respectively (i.e., requiring a regularity cycle and approximately 4 tones). We expect the brain response to follow similar dynamics. All conditions then settle at a low sustained IC, previously shown to mirror sustained response patterning. Therefore, we anticipate these three conditions, REG1, REG3 and REG5, to yield a similar sustained response.

The ideal observer model operates on a symbolic level, and is thus ‘blind’ to low level effects such as adaptation, which are (as discussed above) expected to play a major role in shaping responses to REG1 sequences While REG3 and REG5 contain complex predictable structures that must be discovered on each trial, it is plausible that responses to these stimuli are also influenced by low-level adaptation effects. In REG3, tones are repeated every 150ms, and in REG5 every 250ms. Comparing responses to these patterns at different stages of the stimulus epoch enables a direct comparison of the predictability enhancement and repetition suppression hypotheses.

We analyse EEG responses to these sequences in a time-resolved manner, focusing separately on the onset, sustained, and offset portions of the evoked response to the entire sequence. Onset responses likely reflect initial processing in the auditory cortex, where we anticipate evidence of repetition suppression by comparing REG1 to the other conditions. The sustained response is expected to reflect more complex processing. Our findings reveal that both repetition suppression and repetition enhancement occur over different timescales and that sustained brain responses to simple repetition exhibit qualitatively different effects than for more complex regularities.

## MATERIALS AND METHODS

Electroencephalography (EEG) was used to record the evoked responses to regular (REG) and random (RAND) sequences of tone pips, varying the size of the frequency ‘alphabet’ used. The participants were naïve to the experimental manipulations and were asked to ignore the sounds, watching a silent, subtitled film of their choice throughout.

### Stimuli

Stimuli were 3000-ms sequences consisting of 60 tone pips (50 ms each) with frequencies drawn from a pool of 20 logarithmically spaced values (222 to 2000 Hz; increasing by 12% at each step). Sequences were either regular (REG) or random (RAND) and were generated anew for each trial.

Figure 1a shows schematic spectrograms for each stimulus condition. For the REG conditions, a number alph of frequencies (1, 3 or 5 for REG1, REG3 and REG5 conditions respectively) were randomly chosen from the frequency pool, arranged in random order and then repeated for the duration of the sequence, yielding a regularly repeating pattern. Matched RAND3 and RAND5 sequences were generated by shuffling the REG conditions with alph = 3 and 5. RAND20 was a randomised sequence using the full frequency pool, with equal proportions of all frequencies.

We used an ideal observer model - IDyOM (Pearce, 2005; Pearce et al., 2010; Barascud et al., 2016) to concretize our predictions about the dynamics of predictability within our stimuli. This model tracks the transition probabilities between successive tones. From the model an estimate of the information content of each tone was derived, which can be interpreted as the level of surprise associated with each tone; the lower the value the more predictable it is. This model has previously been shown to closely correspond to the sustained response power and to the latencies at which brain responses diverge from one another (Barascud et al., 2016; Hu et al, 2024). We simulated the detection of sequence predictability for 648 trials in random order, which was equivalent to the order of a true experimental session. The LTM+ model configuration was used, which initiates with an empty model, then learns over the stimulus set, updating the model after each tone. The information content of each tone was averaged over trials of the same condition, and the standard error estimated with bootstrap resampling with 1000 iterations. The resultant plots are provided alongside the stimulus schematics in Figure 1 showing the dynamics of the surprise associated with each tone as regularities become established.

All conditions start with the same information content, then diverge. The information content for REG1 drops substantially from only the second tone, indicating that the model rapidly comes to ‘expect’ tone repetition. REG3 and REG5 show a significant, steep drop in information content at 7 and 9 tones respectively, suggesting that it takes a cycle plus four additional tones for the model to learn the regularity. The information content for REG1 levels off at a value lower than all the conditions, suggesting REG1 is indeed more ‘predictable’ than even the other REG conditions. RAND3, RAND5 and RAND20 show fairly stable information content throughout the sequence, but the information content is lower for successively smaller alphabets, indicating the model successfully detects the stochastic predictability due to restricted alphabet size.

### Procedure

The experiment was conducted in a soundproof booth while subjects watched a muted, subtitled film of their choice.

Trials were presented in random order and each trial consisted of the presentation of a unique sequence of tone-pips, generated according to one of the 6 conditions described above. The inter-stimulus interval (ISI) was jittered between 1100 and 1500 ms. Each condition was repeated 108 times over the entire experiment, for a total of 648 trials overall. Equal proportions of each condition were presented in each of 6 blocks, which lasted for approximately 10 minutes including self-timed breaks. Stimuli were presented binaurally using the Psychophysics Toolbox (Kleiner et al., 2007) for Matlab, using insert earphones (3M E-A-RTONE)

### Recording and Data Preprocessing

EEG signals were recorded using a 64-electrode Biosemi system at a sampling rate of 2048 Hz. Data were analysed using the Fieldtrip toolbox (www.fieldtriptoolbox.org/) (Oostenveld et al., 2010) for Matlab (2017a, MathWorks). Data were split into 5000-ms epochs, with 1000 ms before stimulus onset and after offset. An anti-aliasing 100 Hz low pass filter was applied before down-sampling at 256 Hz. Detrending was applied by fitting a linear trend over 9-second segments, taken from the raw recording, centred on each epoch (de Cheveigné and Arzounian, 2018).

Outlier channels and trials were removed manually using Fieldtrip’s visual artefact rejection tool. Around 95% of trials were retained. Further signal-to-noise improvement was achieved using DSS (de Cheveigné and Parra, 2014) to maximise the reproducibility over all trials, keeping the two components which explained the greatest proportion of the variance over trials. Finally, data were re-referenced to the average over all channels.

Three phases of the evoked response were analysed, the first two represent the sequence-evoked response, and the third the offset-evoked response. (i) The onset-evoked response was considered to be from 0-300 ms post-onset, including the highly stereotyped N1 and P2 waves of the auditory-evoked response. (ii) The sustained response, from 300-3000 ms, where the response power over all channels increases, resembling a DC-shift: polarity reversals generally cease, and this is where the effects of complex sequence regularity are expected to begin. (iii) The offset-evoked response, which resembles the N1-P2 complex, but has been shown to be influenced by the statistics of the preceding sequence (Andreou et al., 2015; Southwell et al, 2018). For ERP plotting and statistical analysis, all evoked responses were low-pass filtered at 30 Hz.

### Sequence-Evoked Responses

Epochs were baseline-corrected relative to the interval 200ms before sequence onset. Epochs were averaged over trials for each condition, yielding the evoked response in each channel. However, for statistical analysis, this evoked response was quantified as the root-mean-square (RMS) over channels, giving a measure of instantaneous evoked power at each time sample.

Statistical analysis was performed separately for the onset and sustained portions of the sequence-evoked response. A two-tail t-test was used to find contiguous clusters of time-points showing pairwise differences between conditions, with the threshold alpha level set at 0.05 (0.025 per tail). Correction for multiple comparisons over time used a non-parametric approach to estimate the significance of the effect in each cluster, using as the test statistic the summed t-values within each cluster. To estimate the null distribution of this statistic, a Monte Carlo permutation approach with 10000 iterations was used (Maris and Oostenveld, 2007). This measure ensures transient differences are less likely to cross the threshold than temporally-extended ones. The existence of sustained differences between conditions will increase the threshold for significance and thus render it less sensitive to any additional effects which occur over short intervals resulting in a smaller summed t-statistic (Maris and Oostenveld, 2007). For this reason, a separate a-priori latency range from 0 ms to 300 ms after stimulus onset was chosen to analyse the onset-evoked response, which is expected to be transient. The sustained response was analysed in the time range 300-3000 ms. Subsequently, bootstrap resampling was used to estimate the mean and the standard error of the responses over subjects, for plotting of the grand averages shown in Figure 2a-e.

**Figure 2:**
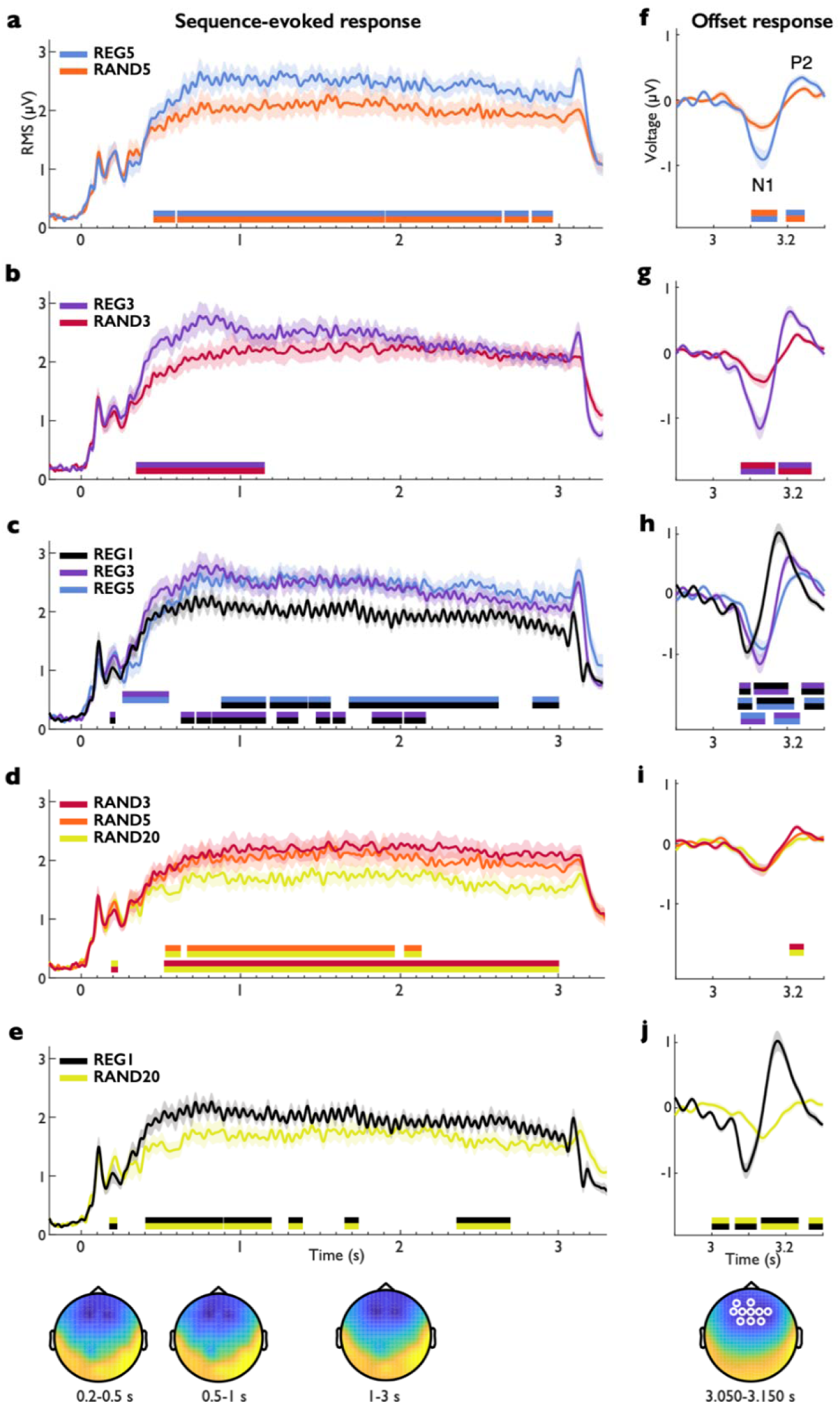
Evoked responses. Bars under plots denote periods of significant difference between conditions; the two colours indicate which pair of conditions are represented. Sustained response shown in a-e; offset response shown in f-j. The lower panel shows topography averaged over all conditions during the specified time ranges. The topography below j indicates 10 channels over which the offset-evoked response is averaged in white.

The following pairs of conditions were contrasted: REG5 vs RAND5; REG3 vs RAND3, REG5 vs REG1; REG5vs REG3; REG3 vs REG1; RAND3 vs RAND5; RAND3 vs RAND20; RAND5 vs RAND20; REG1 vs RAND20. These contrasts correspond to the hypotheses below.

### Offset-Evoked responses

The sequence-evoked response displays a prominent peak at offset in the RMS plot (Figure 2), which was analysed separately as follows. The sequence-evoked epochs were high-pass filtered at 2 Hz and baseline-corrected in the 100-ms window preceding stimulus offset to remove any pre-offset differences in the sequence-evoked response between conditions. These data were then trimmed to epochs with 100 ms before and 300 ms after stimulus offset.

The offset-evoked response was analysed over a subset of channels showing the same response polarity. Taking an RMS over all channels would misrepresent the shape of such a response, as it contains zero-crossings owing to the baseline-correction directly preceding offset. The average signal was taken over the 10 channels showing the strongest negative deflection of the offset response over all subjects and conditions (from 50-150ms post-offset, capturing the initial N1-like deflection). This fronto-central group of channels comprised AFz, AF3, Fz, F1, F2, F3, F4, FCz, FC1, FC2; see topography in Figure 2. The resulting timeseries (one per subject and condition) were compared pairwise, from 3000 ms to 3300 ms, utilising the same cluster-based permutation procedure as for the sequence-evoked response. The latency of the N1 offset peak was computed, for each subject and condition, as the time of the most negative value between 50 and 150 ms post-offset. This was entered into a one-way ANOVA with a factor for each condition.

### Tone repetition rate responses

We calculated EEG response power at the tone repetition rate (20 Hz), corresponding to the IOI (Inter Onset Interval) of 50ms. The power at 20Hz was computed in individual trials between 1000 and 3000 ms, a window chosen to represent the sequence-evoked response once it has stabilised. To obtain a high frequency resolution, these 2.5-second epochs were concatenated to form a single ‘epoch’ per subject and condition. To compensate for different numbers of trials for each subject owing to outlier removal, the concatenated trial was trimmed to 51200 samples for each subject for each condition, to give a frequency resolution of 1/200 Hz. This approach isolates activity with consistent phase across trials because trials were concatenated consistently at the onset of the 10th tone in the sequence. Power over the entire frequency spectrum was calculated using the fast Fourier transform implemented in Fieldtrip’s ft_freqanalysis function, using the multitaper method (Thomson 1982). The resulting power spectra were then RMS-averaged over all channels. The power at 20 Hz was then normalised relative to power of surrounding frequencies, to give a signal-to-noise ratio (SNR). The ‘signal’ was computed as the RMS over the five bins centred on 20Hz. The RMS over the remaining bins between 19 and 21 Hz were taken as ‘noise’; and the SNR computed as SNR= signal/noise. The effect of condition on 20Hz SNR was assessed using a 2-by-2 factorial ANOVA with one factor for alph (3 or 5) and another for regularity (REG or RAND).

### Cycle-rate Responses

It is possible that there is also periodicity in the brain response corresponding to the duration of the repeating cycles in REG3 and 5. Because these sequences were unique on each trial, this effect would most likely be manifest in the induced activity, i.e. non-phase-locked oscillation. However, the inter-trial phase coherence (ITPC) at the cycle rate was also computed. The frequencies of interest correspond to the cycle durations of REG3 (150 ms) and REG5 (250 ms); 6.67 and 4 Hz respectively. The fast Fourier transform was computed for each trial using the multitaper method (Thomson 1982) for frequencies between 1 and 8 Hz. For the power spectral density, the SNR at the frequencies-of-interest was computed as for the tone-locked response, but as the frequency resolution was much lower, only a single bin was taken for the signal and the surrounding 2 bins on each side as noise.

To compute the ITPC for each subject and condition, the complex vectors representing phase and power at the frequency-of-interest in each trial were normalised to unit length, then averaged. The modulus of this averaged vector represents the consistency in phase across trials with a value between 0 and 1 (Lachaux et al. 1999), and the angle of the vector represents the predominant phase. A paired-sample t-test was used to test for differences between REG and RAND, in response power and ITPC, at the cycle rate for alph = 3 and alph = 5.

### Participants

21 normal-hearing young participants took part, with no known neurological or hearing problems. One subject was excluded due to noise during the EEG recording, leaving 20 subjects, 6 male, aged between 20 and 33 (mean 24.4). None of the subjects had participated in similar experiments conducted in the lab. Experimental procedures were approved by the UCL ethics board and written informed consent was obtained from each participant.

## RESULTS

We conducted analyses to address 5 main hypotheses:

1. **REG is associated with an increased sustained response.** As seen previously (Barascud et al, 2016; Southwell et al, 2017; Herrman et al, 2018) we expected that the REG conditions will rise above their matched RAND, diverging at the latency predicted by the ideal observer model (Figure 1; after a cycle plus four tones). A less precise expression of this prediction is that REG3 will diverge from RAND3 before REG5 diverges from RAND5.
2. **RAND sustained response power is inversely related to alphabet size** such that the sustained response magnitude will be RAND3>RAND5>RAND20, mirroring model predictions and replicating the effect seen for larger cycle durations in MEG (Barascud et al. 2016) and EEG (Southwell et al, 2017).
3. **REG is associated with increased offset responses**. An offset response can be seen as a response to the violation of the expectation that a tone will be presented (Southwell &Chait, 2018). This effect has been studied extensively in the context of the auditory omission paradigms (Chennu et al. 2016; Phillips et al. 2016), where unexpected omissions of predictable sounds are associated with a prototypical evoked response. Indeed Southwell & Chait (2018) have shown that offset responses are greater the more predictable the sequence; i.e. greater for REG than for RAND. We expect a similar effect here, potentially modulated by the complexity of the regularity being experienced.
4. **REG1 behaves like REG3 and REG5.** As discussed above, REG1 is the key condition where predictions informed by different literature domains diverge. Based on the sustained response findings it is hypothesised that the sustained response to REG1 will be of similar magnitude as compared to REG3 and REG5, because all are deterministic, and have a similar information content per tone (Figure 1). However it will rise to this level sooner, as the regularity begins from the second tone, as opposed to the fourth or the sixth for REG3 and REG5 respectively. The response will then remain flat until offset. An alternative hypothesis is that the REG1-evoked response will exhibit adaptation, manifest as a decrease of the sustained response throughout the trial.
5. **Regularity is associated with induced power at the cycle rate.** As REG contains a repeated cycle, at a periodicity of 150 or 250 ms for alph = 3 and 5 respectively, it is anticipated that this will be detectible in the brain response as a peak in the frequency spectrum at 6.67 or 4Hz, respectively. This would be expected to be absent for matched RAND, forming the condition against which the presence of a cycle-rate oscillation will be assessed.

Figure 2 shows the average power (RMS over channels) of the evoked responses for each condition. For clarity, only a subset of conditions is shown in each plot, corresponding to key comparisons. Statistical analysis employed a permutation procedure to correct for multiple comparisons over time, to derive the following significant temporal clusters at FWER< 0.05, shown in Figure 2 as horizontal bars below the evoked responses.

All conditions are characterized by a series of onset peaks corresponding to the N1 and P2 time windows, a rise to a plateau (sustained response) for the remainder of the trial, and in some cases a clear offset peak. The offset response is discussed separately below.

### REG3 is associated with a different sustained response pattern to REG5

The effect of sequence regularity, controlled for (long-term) tone probability, is seen when contrasting REG and RAND of the same alphabet size (Figure 2a,b). Based on previous findings with similar stimuli in EEG (Southwell et al., 2017) and MEG (Barascud et al., 2016), it was predicted (Hypothesis #1, above) that the sustained response to REG would be greater than that to RAND throughout the entire sequence, both for alphabet size = 3 and 5. Indeed the sustained response is greater for REG5 than for RAND5 throughout the sequence. REG5 diverges from RAND5 at 453 ms, which is immediately following the fourth tone in the first repeated cycle; exactly as predicted from the ideal observer model (see Figure 1b). The REG5 sustained response remains significantly higher in five clusters covering most of peristimulus time.

From the ideal observer model, the divergence between REG and RAND was expected to be earlier for REG3 than REG5, at around 350 ms which corresponds to 7 tones. The difference between REG3 and RAND3 is significant from 359 ms, which closely matches the prediction. However, REG3 does not entirely behave as hypothesised; whilst the sustained response is initially higher for REG3, from 1051 ms it dropped back to a similar level to the RAND3 response, remaining statistically indistinguishable for the remaining 2 seconds of the stimulus. The response to REG3 rises significantly above REG5 from 258-551 ms (Figure 2c), likely reflecting the earlier discovery of the regularity.

### RAND sustained response power is inversely related to alphabet size

The effect of alphabet size, independent of regular cycle length, is revealed by the comparison between RAND3, RAND5 and RAND20 (Figure 2d; Hypothesis #2). This represents a manipulation of non-deterministic predictability from the most predictable (RAND3) to the least (RAND20). The response to RAND3 is significantly greater than RAND20, from 188-230 ms after stimulus onset, then again from 520 ms until the end of the sustained response. Likewise, the response to RAND5 is higher than to RAND20, but significantly so for less of the sequence (between 527 and 2137 ms). The mean sustained response level appears to be consistently higher for RAND3 than RAND5, though this is not significant.

### REG1 associated with a different sustained response pattern to REG3 and REG5

To investigate the effect of simple tone repetition in regular conditions, REG1 was compared to REG3 and REG5 (Figure 2c; Hypothesis #4). In the onset response, REG3 showed a briefly larger peak than REG1 in the range 180-215 ms. Both REG3 and REG5 evoked a larger sustained response than REG1, despite all being equally predictable (and despite REG1 being associated with the lowest model IC; Figure 1b). This first reached significance at 625 ms for REG3 and 879 ms for REG5. Both contrasts with REG1 did not show stable significance throughout the sequence, although significant clusters were seen for REG5>REG1 until offset. Response power to REG3 dropped throughout the sequence and was no longer significantly different from REG1 after 2164 ms, although the grand average power never drops to that of REG1. This suggests that, unlike the pattern seen for larger cycle durations (e.g. Barascud et al, 2016), shorter cycle durations are associated with a sustained response that progressively reduces in the course of sequence presentation. This will be discussed further below.

REG1 was compared to RAND20 (Figure 2e), the least predictable stimulus, to determine if repetition suppression or repetition enhancement is dominant in the sequence-evoked response. At onset, repetition suppression was seen with RAND20 significantly above REG1 from 176-227 ms. During the sustained response the opposite effect was seen; REG1 response was greater than RAND20 from 402 ms onward, showing a significant effect intermittently throughout the trial.

To summarise, REG sequences evoked a higher sustained response than matched RAND sequences, for alphabet size 3 and 5, although this was only the case for the first second of the sequence for REG3. Within random sequences, decreasing alphabet size was associated with an increased sustained response, thus replicating previous findings (Barascud et al., 2016; Southwell et al., 2017). REG1 did not behave like the other five conditions, exhibiting a nuanced combination of repetition suppression and enhancement. REG1 consistently evoked a lower response throughout the trial compared to REG3 or REG5.

### Offset-evoked EEG responses are modulated by sequence regularity

A different way of probing the brain’s internal model of stimulus regularity is to violate the pattern and observe the deviance response (Southwell & Chait, 2018). In the current experiment there is a form of deviance consisting of the stimulus offset (Andreou et al., 2015); this violates the simple expectation of ongoing sound established over the preceding three seconds (Hypothesis #3).

The offset responses were analysed separately, removing the pre-existing sustained response differences through baseline correction and high-pass filtering. These evoked responses, averaged over a selection of 10 channels to retain the polarity reversals of successive peaks, are shown in Figure 2f-j. All responses resemble the classic N1-P2 responses to sound onset, so are referred to as offset N1 and offset P2 below. One-way ANOVAs were carried out on the peak latency and peak magnitude of the offset N1, for the peaks identified separately for each subject and condition. There was a main effect of condition on peak magnitude (F5,119 = 7.3, p <0.0001) and latency (one-way ANOVA; F5,119 = 7.7, p <0.0001). Post-hoc comparisons used Tukey’s honestly significant difference procedure and are reported below.

### Regularity increased the offset-evoked response for alphabet size = 3 and 5

Figure 2 f-g shows the offset response for REG was substantially larger than for RAND, for alph = 3 and 5. This effect was significant for the offset N1 from 74-168 ms for alph = 3, and from 102-172 ms for alph = 5. For the offset P2, REG was more positive than RAND from 195-245 ms for alph = 5 and from 176-266 ms for alph = 3. REG3 showed a greater negative deflection than REG5 from 74-141 ms, and a correspondingly larger P2 than REG5 from 164-234 ms. These significant effects for REG3 as compared to RAND3 and REG5 are interesting, as the sustained response for REG3 was not different directly preceding offset, suggesting that the offset and sustained responses probe different processes.

### The offset N1 occurred earlier for REG1

Figure 2h shows offset responses to the three REG conditions. For REG1, the offset N1 falls significantly earlier than for the other conditions, with a mean latency of 96 ms, as opposed to over 116 ms for all other conditions. The latency of the N1 did not differ for all other conditions. The N1-latency response for REG1 appears to be smaller in amplitude compared to the N1 responses for REG3 and REG5. However, according to post-hoc comparisons, this difference did not reach statistical significance. REG1 again shows an earlier peak in the P2 window, but it is somewhat larger than for either of the other conditions. As this effect is likely correlated with the earlier N1 latency, the P2 peak was not subject to a separate statistical analysis.

All RAND conditions show the same offset response profile, but greatly attenuated in magnitude as compared to the REG conditions.

### Tone-rate (20Hz) responses are modulated by sequence regularity

The power spectral density (PSD) around 20Hz (corresponding the tone duration of 50ms) is shown in Figure 3a. There is a highly pronounced peak at 20Hz for all conditions. The signal-to-noise ratio (SNR) of the response at 20Hz is shown in Figure 3b representing the tone-locked response; this is much larger for REG1 than for all other conditions. As the four conditions REG3, REG5, RAND3 and RAND5 represent a balanced design, the effects of regularity and alphabet size on the tone-locked response were analysed using a two-way repeated-measures ANOVA, with factors of regularity and alph. This shows a weak main effect of regularity (F1,19 = 5.48, p = 0.030), a weak main effect of alphabet size (F1,19 = 4.69, p = 0.043) and no interaction (F1,19 = 1.636, p = 0.22); the SNR at 20Hz being higher for REG than for RAND, and higher for alph = 3 than alph = 5.

**Figure 3:**
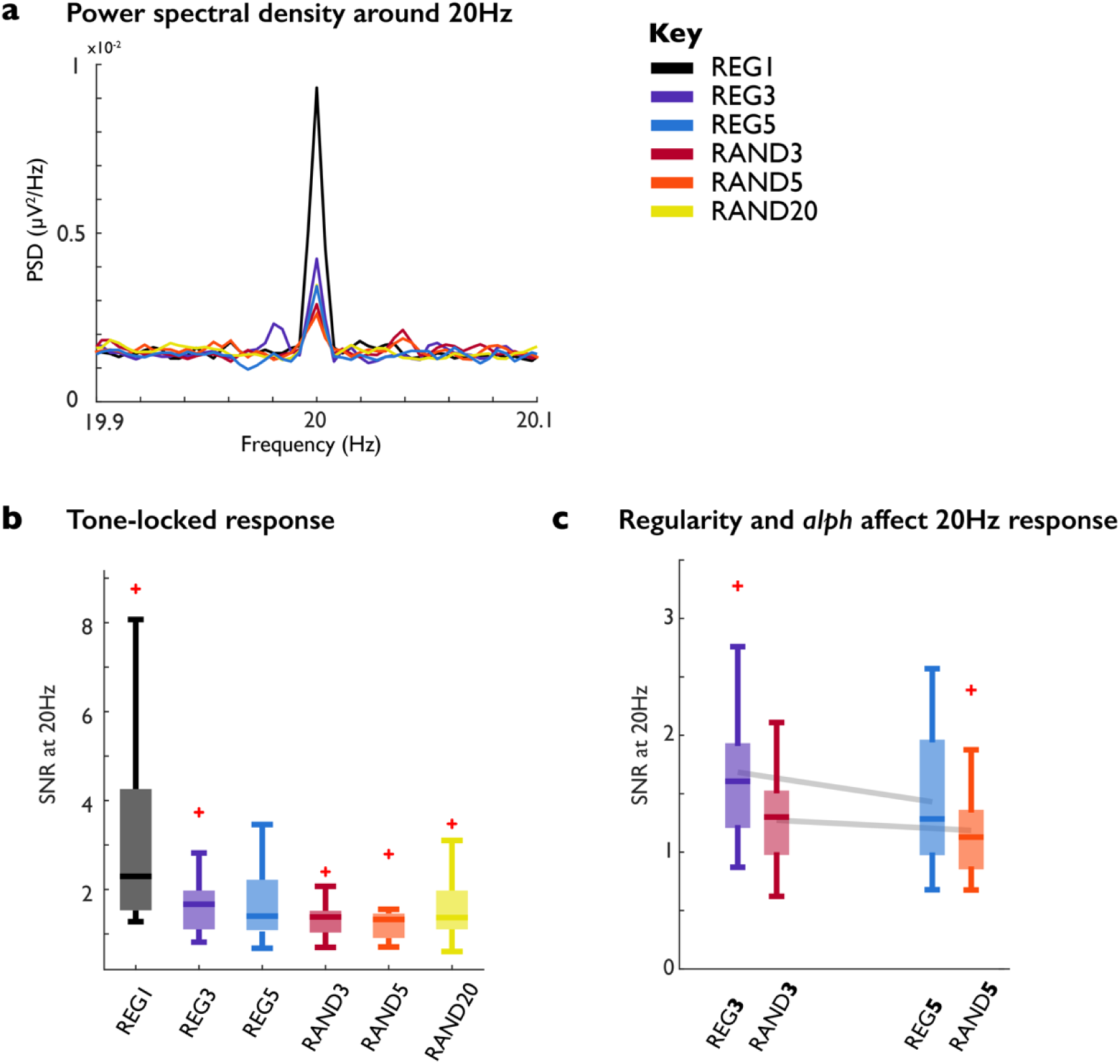
Tone-rate response. Frequency analyses. a: Power spectral density around 20Hz. b: Signal-to-noise ratio at 20Hz for each condition. c: Signal-to-noise ratio at 20Hz for matched REG and RAND conditions. Grey lines connect group means for REG and RAND of different alph. Boxplots show distribution over subjects: median with lower and upper quartiles, whiskers show range. Extreme observations, falling outside of 1.5 times the interquartile range from the start of the whisker, are shown by a red cross.

### Cycle-rate responses are seen in REG3 and REG5, reflecting regular pattern tracking

There is a significant effect of regularity on the induced response at the cycle rate in both REG3 and REG5 as compared to their matched RANDs (Figure 4). REG3 shows a larger response at 6.67 Hz than RAND3 (t19 = 3.03, p = 0.0068), and REG5 shows a larger response at 4 Hz than RAND5 (t19 = 2.40, p = 0.0268). This means that the periodicity of REG is reflected in the response on individual trials, but this is likely to be an idiosyncratic pattern which does not have a consistent phase across trials, because the regularity comprises a different pattern each time. Indeed, the ITPC did not significantly differ between REG and RAND (alph = 3: p = .62, alph = 5: p = 0.51; Figure 4c&d).

**Figure 4:**
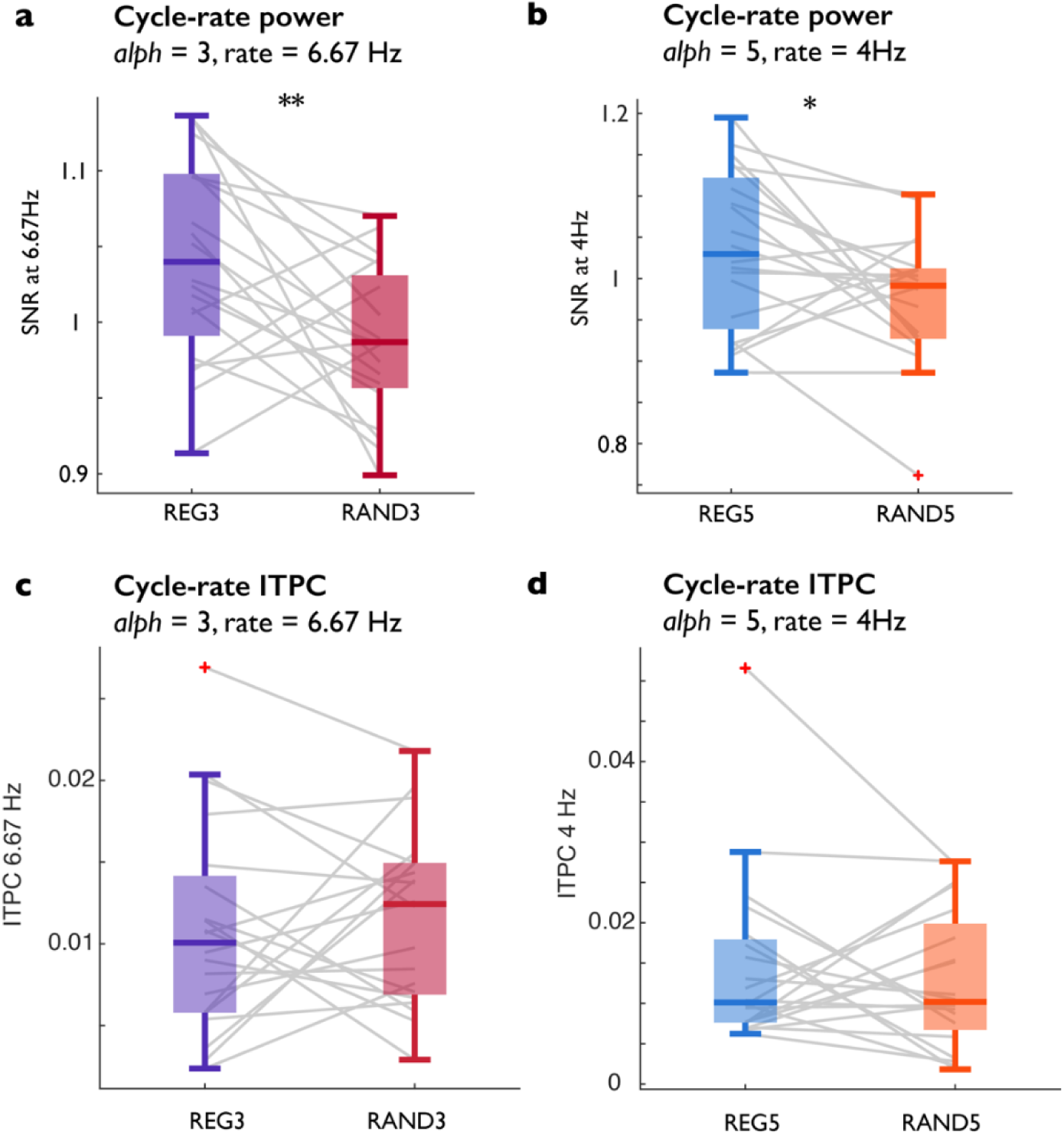
Cycle-rate response. a: Signal-to-noise ratio at cycle rate of 6.67Hz for alph = 3. b: Signal-to-noise ratio at cycle rate of 4Hz for alph = 5. c: ITPC at 6.67Hz for alph = 3. d: ITPC at 4Hz for alph = 5. Boxplots show distribution over subjects: median with lower and upper quartiles, whiskers show range. Extreme observations, falling outside of 1.5 times the interquartile range from the start of the whisker, are shown by a red cross. **p < .01, *p < .05

## DISCUSSION

We examined the influence of sequence structure on various aspects, both evoked and induced, of EEG responses. Our primary objective was to elucidate the way patterns of brain response evolve with the progressive simplification of regularities. We employed three distinct types of regular tone pip sequences: REG5, characterized by cycles of five tones; REG3, featuring cycles of three tones; and REG1, representing tone repetition—a paradigmatic stimulus in the investigation of auditory predictive coding (Baldeweg, 2006). By contrasting responses across different phases of the stimulus—onset, sustained, and offset—we discerned the interplay between repetition suppression and predictability within the EEG response.

### Repetition suppression effects are detectable in onset responses

The long sequences offer the opportunity to observe how the brain response unfolds over several seconds. The manipulations in this experiment affected different phases of the response in different ways. A reduced response was seen in the first 300 ms for REG1 relative to other conditions with a larger alphabet, and for RAND3 relative to RAND20. This could be explained as adaptation to tone frequency, as in all these cases the suppressed condition contains more repetitions over frequencies encountered in the sequence so far. The sub-300-ms timescale of these effects is in line with the timing of repetition suppression observed in multiple paradigms (Budd et al., 1998; e.g. Grill-Spector et al., 2006; Herrmann et al., 2015). This onset effect precedes the rise to the sustained response when, based on previous evidence (Barascud et al, 2016; Hermann et al, 2018; Hu et al, 2024; Zhao et al, 2024), the representation of predictability influences the response magnitude.

### The sustained response is transient for increasingly simple sequences

As demonstrated previously for larger alphabets (Barascud et al, 2016) we observed that the more restricted the alphabet in random sequences, the greater the sustained response magnitude, which is in line with the decreased information content of each tone in RAND with a restricted alphabet (Figure 1b). Although this is surprising from the standpoint of neuronal adaptation (which would have been expected to yield the opposite pattern), there is accumulating evidence that the sustained response encodes this aspect of sequence predictability (Zhao et al, 2024; Barascud et al, 2016).

Similarly, regularity boosted the sustained response over random sequences of the same alphabet size. The timing of the initial divergence between REG and RAND was related to alphabet size as predicted by the model (Figure 1). REG5 showed a sustained difference from RAND5 for the duration of the epoch consistent with what was seen for longer patterns in previous work and hypothesized to reflect the encoding of the predictability of unfolding patterns (Barascud et al, 2016; Zhao et al, 2024; Hu et al, 2024).

However, for REG3 this effect was transient before the evoked power dropped back to the level of RAND3, following around 7 repetition cycles. Notably the response pattern suggests a stable RAND response and a gradual decrease in the responses to REG3, despite both being matched in frequency content.

There are two possible explanations for the observed effects: one related to an interplay between predictability and adaptation, and another related to an inherent process associated with overlearned patterns.

In the first explanation, the initial dominance of predictability leads to higher responses to REG compared to matched RAND sequences. However, as the trial progresses, adaptation sets in, gradually reducing responses to REG stimuli. RAND sequences may not exhibit a similar effect due to their random structure, which may result in less overall adaptation (e.g. as shown by Costa-Faidella et al, 2014).

According to the second explanation, the effects observed for REG3 may reflect a shift in the system’s operation for overlearned patterns, leading to changes in how predictability is encoded. The sustained response, which involves a network comprising the hippocampus, inferior frontal gyrus (IFG), and auditory cortex (Barascud et al, 2016; Hu et al, 2024), conveys predictions from higher-level regions (IFG) to the auditory cortex. Once regularities are overlearned, resulting in near-zero prediction errors, the system may shift to a different dynamic, leading to the abolishment of the sustained response. Longer patterns, such as REG5 or REG10, may also exhibit similar effects if presented for a sufficiently long duration. However, previous experiments have typically presented a maximum of 5 cycles, potentially obscuring these effects.

Importantly the effects of suppression appeared to be specific to the sustained response – other measures of sensitivity to regularity (i.e. the difference between matched REG and RAND sequences) including the offset response, the 20Hz response, and the cycle-locked response continued to indicate significant differences between REG3 and RAND3.

The two explanations above may also underly the distinct pattern exhibited by the REG1-evoked sustained response, which was lower than for REG3 and REG5, despite being as predictable. REG1 represents the most predictable stimulus, which also theoretically leads to the greatest degree of adaptation/repetition suppression in units selective for the repeated frequency. What is clear is that the responses to this stimulus cannot solely be described by repetition suppression, as demonstrated most strongly by the comparison with RAND20. The response was still higher than to RAND20, both during the sustained response and at offset.

To summarise, predictability in two forms is associated with an increase in brain response; whether the predictability arises from a deterministic order of frequencies being repeated (REG) or from a restricted alphabet size (RAND3 as compared to RAND5 and RAND20). However, this was not universally the case; notably the response to REG1 behaved differently from the rest in several respects. The sustained response to REG1 never reached as high as for REG3 and REG5, despite being equally predictable. Yet, the response to maximally-repetitive REG1 was still higher than for the least adapting stimulus, RAND20. This suggests that opposing effects of repetition suppression and regularity enhancement influence the sustained response.

### Repetition suppression vs. repetition enhancement

Predictability-related suppression of measured responses can result from a combination of different underlying processes, whether neuronal refractoriness, prediction error suppression or sharpening of representations (Desimone, 1996; Budd et al., 1998; Grill-Spector et al., 2006; Kok et al., 2012). Even at the neuronal level within a given cortical area, adaptation to repetitive stimuli affects excitatory and inhibitory inputs at different timescales (Solomon and Kohn, 2014; Chen et al., 2015), and has complex downstream effects (Solomon and Kohn, 2014). At the population level, different timescales of repetition effects correspond to different cognitive phenomena. Expectation suppression and repetition suppression to sound have been demonstrated to have distinct temporal profiles (Todorovic and de Lange, 2012), with repetition affecting earlier response components (between 40 and 60 ms) than expectation (from 100 to 200 ms). This effect was generalised to the visual modality in fMRI (Grotheer and Kovács, 2015) and EEG (Feuerriegel et al., 2017). These findings support a two-stage account of repetition suppression, incorporating a hierarchically-higher stage of expectation suppression following an earlier primary-sensory stage of stimulus-specific adaptation (Grotheer and Kovács, 2016). The divergent effects on the sustained response of simple repetition (REG1) and complex regularities (the other 5 conditions) seen here supports a similar model.

Although the present experiment involved passive listening in naïve subjects, so any expectations formed on the basis of regularities are most likely implicit, a similar multi-stage model could account for at least some of the results here. However, distinct from the cited findings, this study revealed multiple instances of enhancement according to predictability, even for the simplest model of predictability engendered by the repetition in REG1. Repetition enhancement, although less investigated than repetition suppression, has been reported in multiple domains (Segaert et al., 2013). It is most often observed in explicit paradigms requiring behavioural responses such as priming (e.g. Petit et al., 2006; Henson et al., 2008) but it has been reported in responses to passively-presented stimulation (Stefanics et al., 2018). Of most relevance to the present study, concomitant repetition suppression and repetition enhancement effects were observed in an MEG study of passively presented auditory roving-standard sequences (Recasens et al., 2015). Repetition suppression occurred during the N1m, and was localised to temporal cortical sources. Later repetition enhancement was observed between 200 and 300 ms during a sustained potential with greater field strength than the N1m, with generators in the same temporal areas but also in inferior frontal gyrus. It is tempting to relate this to the current EEG sustained response displaying repetition enhancement when comparing REG1 and RAND20.

Predictive coding provides two neuronal mechanisms for response enhancement (precision and prediction), and one mechanism for response suppression (reduced prediction error), consistent with the processing of a predictable stimulus stream. The sequences used here represented a constant level of predictability throughout each trial, such that by the first half-second or so of each stimulus, it is possible to form an expectation of both the identity of the next tone and the precision of this estimate, given the context provided by the previous tones. The neuronal mechanism for precision signalling in predictive coding is postsynaptic gain on the superficial pyramidal prediction-error units, leading to ascending prediction errors up-weighted by (expected) precision (Friston, 2005; Feldman and Friston, 2010). Such increased superficial pyramidal activity plausibly forms a large contribution to measured EEG signals (Murakami & Okada 2006). Alternatively, sustained responses may indicate an increase in inhibitory activity among neuronal units conveying low information content. This notion aligns with previous research, albeit with simpler stimuli, demonstrating heightened inhibitory activity in the presence of predictable information (Natan et al., 2015, 2017; Schulz et al., 2021; Richter and Gjorgjieva, 2022; Yarden et al., 2022). The emphasis on inhibition, rather than excitation, in modulating responses to predictable sensory stimuli finds support from indirect evidence, including dynamic causal modelling (Lecaignard et al., 2022) and behavioural studies. Predictable patterns are more easily disregarded rather than capturing attention (Southwell et al., 2017) and are associated with diminished arousal (Milne et al., 2021). Furthermore, recent findings indicating a decline in the sustained response with aging (Hermann et al., 2023) align with the inhibition hypothesis, given the documented reduction in inhibition within the aging brain (Caspary et al., 2013; Hughes et al., 2010; Rozycka & Liguz-Lecznar, 2017).

### Tone-locked and cycle-locked responses reflect sequence predictability

We observed systematic effects on the 20Hz response, reflecting tone-rate activity. Specifically, REG1 elicited the highest power, followed by REG3 and then REG5. Additionally, there was an interaction between alphabet size and predictability, indicating higher responses for regular sequences (REG) compared to matched random sequences (RAND), particularly for smaller alphabet sizes.

The observed pattern of increased responses to REG compared to RAND aligns with the precision weighting mechanism discussed earlier. However, it may seem contradictory to recent findings showing reduced responses to individual tones within REG patterns (specifically between about 100 - 200 ms post onset; Hu et al., 2024). It is possible that the discrepancy is rooted in the difference in tone rates, with responses to faster tones behaving differently than those to slower tones examined in Hu et al. (2024). However, we think a more likely explanation is that the 20 Hz response does not reflect the same processes as the evoked responses analysed in Hu et al. (2024) but rather echoes increased phase locking, indicating more accurate timing of responses in predictable sequences. This interpretation aligns with the notion that the 20 Hz response may serve as a marker of temporal synchronization, emphasizing the brain’s ability to accurately anticipate and time its responses to predictable stimuli. Therefore, while both the 20 Hz response and tone-evoked responses (as measured in Hu et al., 2024; but unavailable here due to the rapid presentation rate) may reflect aspects of predictive processing, they likely represent distinct facets of neural activity involved in processing auditory regularities.

Furthermore, we observed effects at cycle duration, consisting with automatic tracking of the regular structure. Specifically, REG3 was associated with increased activity at 6.6Hz, and REG5 was associated with increased activity at 4 Hz—values corresponding to the pattern duration, indicating tracking of this periodicity in the unfolding sequence. A similar phenomenon has been previously demonstrated in studies of statistical learning (Batterink et al., 2017; Ding et al., 2016) where exposure to a structured stream of repeating nonsense words is associated with EEG entrainment at the repetition frequency. The fact that these effects were only seen in cycle rate power suggests that they are unlikely to reflect simple responsivity or adaptation effects, as they were not observed in inter-trial phase coherence (ITPC). Herrmann and Johnsrude (2018), using REG10 patterns, similarly did not observe effects in ITPC between REG and random sequences, consistent with our findings. However, they did not examine a phase-insensitive measure of the cycle-rate response, such as induced response power, as utilized here.

Both effects—increased responses in REG over RAND (and larger for REG3 than REG5) at the cycle rate and tone-rate—appear functionally distinct from the sustained response, providing further insights into the neural mechanisms underlying processing of predictable sequences.

Overall, our results demonstrate that subtle interactions between enhancement and suppression of auditory responses are involved in automatic tracking of stimulus statistics, implicating multiple, parallel processes.

## DATA AVAILABILITY

The data reported in this manuscript will be made available after publication at DOI: 10.5522/04/26325733

## ACKNOWLEDGMENTS

This work was supported by a BBSRC project grant to MC and a Brain Trust PhD studentship to RS. The funders had no role in study design, data collection and analysis, decision to publish or preparation of the manuscript.

## AUTHOR CONTRIBUTIONS

MC and RS Conceived and designed research; RS and CT performed experiments and analyzed data; MC and RS interpreted results of experiments; RS prepared figures, drafted manuscript; MC, RS and CT edited and revised manuscript, approved final version of manuscript.

## Notes

### Competing Interest Statement

The authors have declared no competing interest.

